# Sublethal β-lactam antibiotics induce PhpP phosphatase expression and StkP kinase phosphorylation in PBP-independent β-lactam antibiotic resistance of *Streptococcus pneumoniae*

**DOI:** 10.1101/342188

**Authors:** Yan-Ying Huang, Yan-Hong Sun, Peng Du, Xiao-Xiang Liu, Jie Yan, Ai-Hua Sun

**Author notes:** Authors contributed equally to this work (Yan-Ying Huang, Yan-Hong Sun). Corresponding author: Ai-Hua Sun, Faculty of Basic Medicine, Hangzhou Medical College, Hangzhou, Zhejiang 310053, P.R. China. E-mail addresses: Yan-Ying Huang, Yan-Hong Sun, Peng Du, Xiao-Xiang Liu, Jie Yan, Ai-Hua Sun.

## Abstract

StkP and PhpP of *Streptococcus pneumoniae* have been confirmed to compose a signaling couple, in which the former is a serine/threonine (Ser/Thr) kinase while the latter was annotated as a phosphotase. StkP has been reported to be involved in penicillin-binding protein (PBP)-independent penicillin resistance of *S. pneumoniae*. However, the enzymatic characterization of PhpP and the role of PhpP in StkP-PhpP couple remain poorly understood. Here we showed that 1/4 minimal inhibitory concentration (MIC) of penicillin (PCN) or cefotaxime (CTX), the representatives of β-lactam antibiotics, could induce the expression of *stkP* and *phpP* genes and phosphorylation of StkP in PCN/CTX-sensitive strain ATCC6306 and three isolates of *S. pneumoniae* (MICs: 0.02-0.5 μg/ml). The product of *phpP* gene hydrolyzed PP2C type Ser/Thr phosphotase-specific RRA(pT)VA phosphopeptide substrate with the Km and Kcat values of 277.35 μmol/L and 0.71 S^−1^, and the hydrolytic activity was blocked by sodium fluoride, a PP2C type Ser/Thr phosphatase inhibitor. The phosphorylation levels of StkP in the four *phpP* gene-knockout (Δ*phpP*) mutants were significantly higher than that in the wild-type strains. In particular, the MICs of PCN and CTX against the Δ*phpP* mutants were significantly elevated as 4-16 μg/ml. Therefore, our findings confirmed that sublethal PCN and CTX act as environmental inducers to cause the increase of *phpP* and *stkP* gene expression and StkP phosphorylation. PhpP is a PP2C type Ser/Thr protein phosphatase responsible for dephosphorylation of StkP. Knockout of the *phpP* gene results in a high level of StkP phosphorylation and PBP-independent PCN/CTX resistance of *S. pneumoniae*.

**Importance:** *Streptococcus pneumoniae* is a common pathogen in human populations in many countries and areas due to the prevalence of β-lactam antibiotic-resistant pneumococcal strains. Production of β-lactamases and mutation of penicillin-binding proteins (PBP) have been considered as the major β-lactam antibiotic-resistant mechanisms in bacteria, but *S. pneumoniae* has not been confirmed to produce any β-lactamases and many pneumococcal strains present PBP mutation-independent β-lactam antibiotic resistance. StkP is a Ser/Thr kinase of *S. pneumoniae* to compose a signal-couple with PhpP protein. The present study demonstrated that the PhpP is a PP2C-type phosphotase for dephosphorylation of StkP and the sublethal penicillin (PCN) or cefotaxime (CTX) acted as environmental signal molecules to induce the expression of PhpP. The knockout of PhpP-encoding gene caused the PCN/CTX resistance generation of PCN/CTX-sensitive pneumococcal strains. All the data indicate that StkP-PhpP couple of *S. pneumoniae* is involved in PBP mutation-independent β-lactam antibiotic resistance by phosphorylation of StkP.

## Introduction

*Streptococcus pneumoniae* is a major causative agent of bacterial pneumonia and tympanitis in children [1-3]. More importantly, in the recent years, *S. pneumoniae*-infected meningitis cases with high fatality have been frequently reported in many countries and areas [4-8]. Therefore, *S. pneumoniae* is a common pathogen for human beings with global importance.

β-lactam antibiotics are the first choice in clinic to cure *S. pneumoniae*-infected patients [9]. However, in the recent years, β-lactam antibiotic-resistance of *S. pneumoniae* isolates from patients is continuously increased and the antimicrobial-resistant *S. pneumoniae* strains became more epidemic in many countries and areas [10-15], which has been considered as the major reason for increased incidence of *S. pneumoniae*-infected diseases [15,16]. Bacterial β-lactamases have been confirmed to play a key role in generation of β-lactam antibiotic resistance in many bacteria including *S. pneumoniae* [17]. Mutation of penicillin-binding proteins (PBP), the receptors of β-lactam antibiotics located on surface of bacteria, has been reported as the major β-lactam antibiotic resistant mechanism in bacteria [18,19]. However, recent studies found that some of the *S. pneumoniae* strains had no PBP mutation but presented β-lactam antibiotic resistance [20-22], indicating that *S. pneumoniae* may have a PBP mutation-independent mechanism of β-lactam antibiotic resistance.

StkP is a sequence-conserved eukaryotic-type serine/threonine (Ser/Thr) kinase (STK) of *S. pneumoniae* that has been confirmed to be involved in PBP mutation-independent penicillin resistance [22]. In the chromosomal DNA of *S. pneumoniae*, StkP-encoding gene (*stkP*) and *phpP* gene compose a *stkP*-*phpP* operon and the product of *phpP* gene is annotated as a putative phosphatase [23,24]. A previous study demonstrated that the PhpP and StkP of *S. pneumoniae* composed a StkP-PhpP signaling couple [25]. It has been reported that both prokaryotic and eukaryotic STKs are activated through phosphorylation at Ser/Thr sites and some certain protein phosphatases can inactivate STKs by hydrolysis of phosphoryl groups at the Ser/Thr residual sites in STKs [26]. In particular, a previous study revealed that penicillin (PCN) could cause the gene expression profile change of *S. pneumoniae* [27]. Therefore, we presume that β-lactam antibiotics may act as environmental inducers to cause the change of PhpP and StkP expression and StkP dephosphorylation of *S. pneumoniae* as well as the PhpP may be involved in the StkP-associated PBP mutation-independent penicillin resistance by dephosphorylation of StkP.

In the present study, we used PCN and cefotaxime (CTX) as the representatives of β-lactam antibiotics to detect their induction of *phpP* and *stkP* gene expression and then identified the product of *phpP* gene as a PP2C type Ser/Thr protein phosphatase by virtue of its ability to hydrolyze Ser/Thr phosphatase-specific substrates. Moreover, the *phpP* genes of *S. pneumoniae* strains were inactivated to determine the role of PhpP in dephosphorylation of StkP *in vivo* and the change of β-lactam antibiotic resistance. The results of this study confirmed that the product of *S. pneumoniae phpP* gene is a Ser/Thr protein phosphatase that involved in β-lactam antibiotic resistance-associated StkP-PhpP signaling couple by StkP dephosphorylation during induction of sublethal PNC and CTX.

## Materials and Methods

### Bacterial strains and culture

*S. pneumoniae* ATCC6306 and three β-lactam antibiotic-sensitive *S. pneumoniae* isolates (No.: SP5, SP9 and SP14, belonging to serotype 3, 19F and 19A) from pneumonia children were kindly provided by the Department of Medical Microbiology and Parasitology, Zhejiang University School of Medicine. All the strains were cultured with Colombia blood agar (bioMerieux, France) or 0.5% yeast extract-containing Todd-Hewitt (TH) broth (Sigma, USA) at 37°C [28]. Besides, *Escherichia coli* EL21DE3 (Novagen, USA) was cultured in Luria-Bertani (LB) medium (Oxoid, England) at 37°C.

### Animal

New Zealand white rabbits (3.0 to 3.5 kg per animal) were provided by the Laboratory Animal Center of Hangzhou Medical College (Certificate No.: SCXK[zhe]2012-0173). All the animal experimental protocols were approved by the Ethics Committee for Animal Experiment of Hangzhou Medical College.

### Drug susceptibility test

Susceptibility of each of the four *S. pneumoniae* strains to PCN or CTX was detected by E-test (bioMerieux) according to the manufacturer’s instruction. The minimal inhibitory concentrations (MIC) against *S. pneumoniae* strains, ≤ 2 or ≥ 8 μg/ml of PCN (Sigma) and ≤ 1 or ≥ 4 μg/ml of CTX (Sigma), were considered to be sensitive or resistant [29].

### Detection of sublethal PCN- and CTX-induced expression of *phpP* and *stkP* genes

Each of the four *S. pneumoniae* strains was inoculated into TH broth for a 200 rpm shaking incubation at 37°C. When the value of optical density at 600 nm _(OD600)_ of pneumococcal culture turbidity reached 0.5, 1/4 MIC PCN or CTX was added and then incubated for 0.5, 1, 2, 4, 8 or 12 h as above. After centrifugation and washing with phosphate buffered saline (PBS), total RNA of each of *S. pneumoniae* strains was extracted using a TRIzol^®^ Max^TM^ Bacterial RNA Isolation kit (Invitrogen) plus a gDNA Eraser Kit (TaKaRa, China) and then quantified by ultraviolet spectrophotometry [30]. Subsequently, cDNA from each of the total RNAs was synthesized using a PrimeScript^TM^ RT Reagent Kit (TaKaRa). Using each of the cDNAs as template, the *phpP*- or *stkP-*mRNA level was assessed by real-time fluorescence quantitative reverse transcription polymerase chain reaction (qRT-PCR) with the primers P-1F/P-1R or S-1F/S-1R (Table 1) using a SYBR^®^ Premix Ex-Taq^TM^ Kit (TaKaRa) in an ABI 7500 Real-Time PCR System (ABI, USA). The primers used were designed using Primer Premier 5.0 software according to the *phpP* or *stkP* gene sequences in GenBank (accession No.: NC_003098 and NC_003028). In the qRT-PCR, 16S rRNA gene of *S. pneumoniae* was used as the internal reference [31], while the PCN- or CTX-untreated *S. pneumoniae* strains were used as the controls. The obtained qRT-PCR data were analyzed using the ΔΔCt model and randomization test in REST 2005 software [32].

**Table 1.**
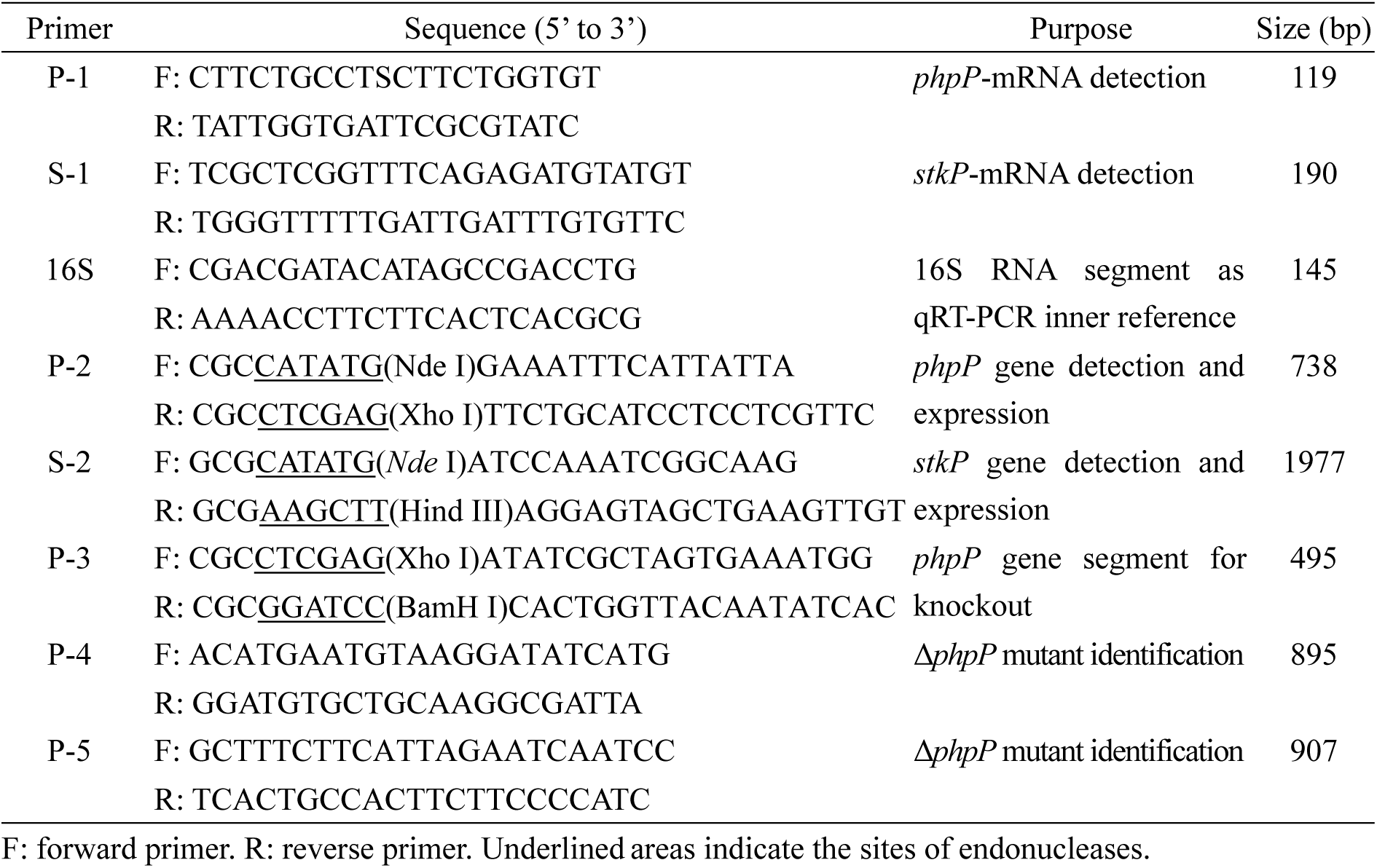
Sequences of primers used in this study.

### Amplification and sequencing of *PhpP* and *stkP* gene segments

Genomic DNA of each of the four *S. pneumoniae* strains was extracted using a Bacterial Genomic DNA Preparation Kit (Axygen). By using a High Fidelity PCR Kit (TaKaRa), the entire *phpP* or *stkP* gene segments were amplified from the DNA templates by PCR using the primers P-2F/P-2R or S-2F/S-2R (Table 1). The PCR products were examined by 1.5% ethidium bromide-stained agarose gel electrophoresis and then cloned into pMD19-T plasmid using a T-A Cloning Kit (TaKaRa) to form recombinant pMD19-T*^phpP^* and pMD19-T*^stkp^* plasmids for sequencing by Invitrogen Co. in Shanghai of China.

### Bioinformatic analysis of *phpP* and *stkP* genes

Since the nucleotide and amino acid sequence identities of *phpP* and *stkP* genes from the four *S. pneumoniae* strains were as high as 98.7%-100%, the *phpP* and *stkP* genes of *S. pneumoniae* strain ATCC6306 were analyzed using TMHMM and NCBI Database Conserved Domain Database (CDD) software [33].

### Generation of prokaryotic expression systems of *phpP* and *stkP* genes

The pMD19-T*^phpP^* or pMD19-T*^stkP^* plasmid from *S. pneumoniae* strain ATCC6306 and pET42a vector (Novagen) were digested with both Nde I and Xho I or Nde I and Hind III (TaKaRa). The recovered *phpP* or *stkP* gene segment was linked with the linearized pET42a using T4 DNA ligase (TaKaRa) and then transformed into *E. coli* BL21DE3 by CaCl_2_ transformation method to form *E. coli* BL21DE3^pET42a-*phpP*^ or *E. coli* BL21DE3^pET42a-*stkP*^. The engineered strains were cultured in LB medium containing 50 μg/ml kanamycin (Sigma) and the pET42a-*phpP* and pET42a-*stkP* were extracted from the strains using a Plasmid Extraction Kit (Axygen) for sequencing again.

### Expression of *phpP* and *stkP* genes and extraction of expressed products

The engineered strains, *E. coli* BL21DE3 ^pET42a-*phpP*^ and *E. coli* BL21DE3^pET42a-*stkP*^, were cultured in kanamycin-containing LB liquid medium to express the target recombinant proteins (rPhpP and rStkP) under induction of 0.5 mM isopropy-β-D-thiogalactoside (IPTG, Sigma). After ultrasonic breakage on ice and a 13,800×g centrifugation for 10 min (4°C), the supernatants of cultures were collected to extract soluble rPhpP and rStkP using a Ni-NTA affinity chromatographic column (BioColor, China). The extracted rPhpP or rStkP was quantified using a BCA Protein Assay Kit (Thermo Scientific, USA). Both the expressed and extracted rPhpP and rStkP were examined by sodium dodecyl sulfate polyacrylamide gel electropheresis (SDS-PAGE) plus a Gel Image Analyzer (Bio-Rad, USA).

### Preparation of rPhpP-IgG and rStkP-IgG

New Zealand rabbits were intradermally immunized on days 1, 14, 21 and 28 with Freund’s adjuvant-mixed 2 mg rPhpP or rStkP per animal. Fifteen days after the last immunization, the sera were collected to separate rStkP-IgG or rPhpP-IgG by ammonium sulfate precipitation plus a DEAE-52 column chromatography using 10 mM phosphate buffer (pH 7.4) for elution [34]. The titer of rPhpP-IgG or rStkP-IgG binding to rStkP or rPhpP was detected by immunodiffusion test.

### Phosphatase activity assays

The enzymatic activity of rPhpP was detected using a pNPP-Phosphate Assay Kit (BioAssay Systems, USA) and a PP2C-specific Ser/Thr Phosphatase Assay Kit (Promega, USA) [35,36]. Briefly, 0.5, 1, 2.5, 5, 10 or 20 μg rPhpP was mixed with 500 nM para-nitrophenyl phosphate (p-NPP), an universal Ser/Thr phosphatase substrate, in 100 μl reaction buffer. After a 30-min incubation at 37°C, the OD_405_ values reflecting p-NPP hydrolysis were detected using type M5 spectrophotometer (Bio-Rad). On the other hand, 0.5, 1, 2.5, 5, 10 or 20 μg rPhpP was mixed with 100 μM RRA(pT)VA phosphopeptide, a PP2C type Ser/Thr protein phosphatase-specific substrate, in 100 μl reaction buffer. After incubation as above, the OD_600_ values reflecting RRA(pT)VA hydrolysis were detected by spectrophotometry.

### Phosphatase inhibition test

Okadaic acid (OA) is an inhibitor of PP1, PP2A and PP2B type Ser/Thr phosphatases while sodium fluoride (NaF) is a PP2C type Ser/Thr phosphatase inhibitor [37,38]. Briefly, 5 μg rPhpP was mixed with 0.1, 0.5, 1, 5 or 10 μM OA (Sigma) or 0.1, 0.5, 1, 5 or 10 mM NaF (Sigma) in 100 μl reaction buffer and then incubated at 37°C for 30 min. The activity of OA- or NaF-treated rPhpP to hydrolyze RRA(pT)VA substrate was detected as described above.

### Determination of Km and Kcat values

According to the results of phosphatase activity assays, 5 μg rPhpP was mixed with 50, 100, 150, 200 or 250 μM RRA(pT)VA substrate and then detected by spectrophotometry as described above. According to the OD_600_ values reflecting RRA(pT)VA hydrolysis and free phosphate concentration standard curve, the Km and Kcat values of rPhpP hydrolyzing the substrate were calculated by double reciprocal Lineweaver-Burk plot [39].

### Generation and identification of *phpP* gene-knockout mutant

pEVP3 plasmid has been used to generate the target gene-knockout mutant of *S. pneumoniae* [40,41]. Briefly, a 495-bp *phpP* gene segment (*phpP*-495) from *S. pneumoniae* strain ATCC6306 was amplified using a High Fidelity PCR Kit (TaKaRa) with the primers P-3F/P-3R and then cloned into pMD19-T to form pMD19-T*^phpP^*^−495^ for sequencing as above. The pMD19-T*^phpP^*^−495^ and pEVP3 plasmid were digested with both Xho I and BamH I (TaKaRa). The recovered *phpP*-495 segment was linked with the linearized pEVP3 using T4 DNA ligase (TaKaRa) to form suicide plasmid pEVP3*^phpP^*^495^ for sequencing again. Each of the four *S. pneumoniae* strains was inoculated in TH broth containing 0.01% CaCl_2_ and 0.2% BSA (Sigma) for a 250-rpm incubation at 37°C to the OD_600_ value as 0.25 and then collected by centrifugation. The competent pneumococcal cells were suspended in 200 µl TH broth containing 0.01% CaCl_2_, 0.2% BSA and 200 ng/ml competence stimulation peptide (CSP, AnaSpec, USA) and then added with 200 ng pEVP3*^phpP^*^−495^ for transformation. The mixtures were smeared on 5% sheet blood Columbia agar (bioMerieux) plates containing 2.5 µg/ml chloromycin (CM, Sigma) for a 48-h incubation at 37°C in a 5% CO_2_ atmosphere to obtain *phpP* gene-knockout colonies [42]. The strategy for generating *phpP* gene-knockout mutants (Δ*phpP*-6306, Δ*phpP*-SP5, Δ*phpP*-SP9 and Δ*phpP*-SP14) is summarized in Fig. 1.

**Fig. 1.**
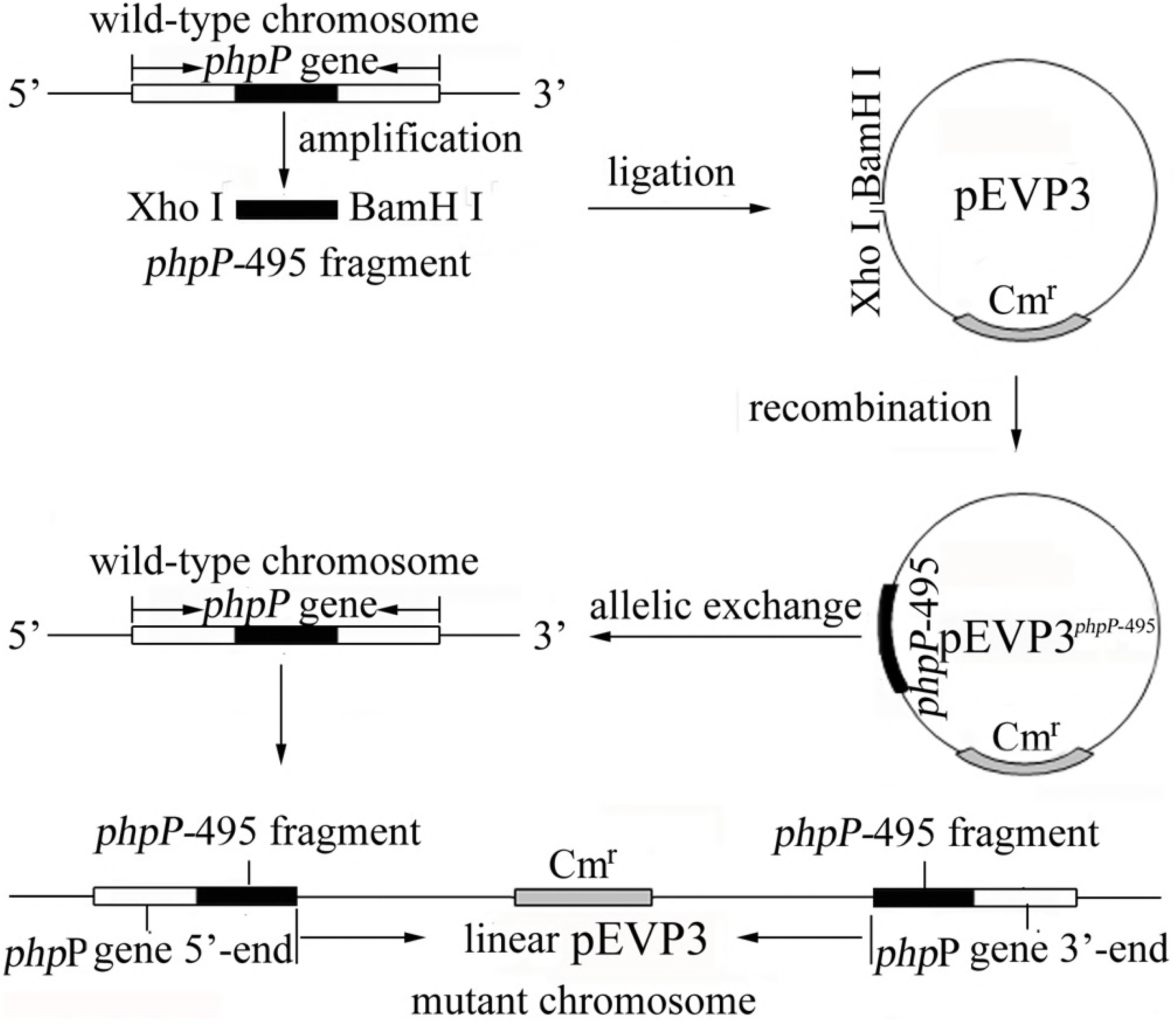
Strategy for generation of *S. pneumoniae* Δ*phpP* mutant. See section of materials and methods in text for details.

### Identification of *phpP* gene-knockout mutants

Growth of the Δ*phpP*-6306, Δ*phpP*-SP5, Δ*phpP*-SP9 or Δ*phpP*-SP14 mutant was assessed by spectrophotometry. The *phpP* gene knockout in the Δ*phpP* mutants was determined by PCR using the primers P-4F/P-4R and P-5F/P-5R (Table 1) and sequencing of the 5’-*phpP*-pEVP3 and pEVP3-3’-*phpP* segment amplification products. Using 1:200 diluted rabbit anti-rPhpP-IgG as the primary antibody and HRP-conjugated goat anti-rabbit-IgG (Abcam, USA) as the secondary antibody, Western blot assay was performed to detect the PhpP form the Δ*phpP* mutants, in which the wild-type strains were used as the controls.

### Detection of sublethal PCN- or CTX-induced StkP phosphorylation

The Δ*phpP* mutants and their wild-type strains were treated with 1/4 MIC PCN or CTX for 1, 2, 4 or 8 h at 37°C. After centrifugation and washing with PBS, the pneumococcal pellets were suspended in deionized water and then ultrasonically broken on ice. The lysates were centrifuged at 14,000×g for 10 min (4°C) and then the supernatants were collected to detect protein concentrations using a BCA Protein Assay Kit (Thermo Scientific). 200 µg of each of the total pneumococcal proteins was mixed with 20 µg rabbit anti-rStkP-IgG for a 2-h incubation in a 90-rpm rotator (4°C). The mixture was added with 600 µg protein-A-coated agarose beads (Millipore, USA), followed by a 60-min incubation as above. After a 14,000×g centrifugation for 5 min and washing thoroughly with PBS, the precipitated beads were suspended in 200 µl 50 mM Tris-HCl buffer (TB, pH7.5) and then mixed with 200 µl 2M NaOH solution for a 30-min water-bath at 65°C to release phosphate groups from the captured StkP according to the instruction of Phosphoprotein Phosphate Detection Kit (Sangon Biotech, Canada). The mixture was neutralized with 200 µl 4.7 M HCl solution and then added with 200 µl detection buffer for a 30-min incubation at room temperature. The OD_620_ value reflecting phosphate group concentration released from IgG-captured StkP were detected using a spectrophotometer (Molecular Devices, USA) [43]. In the detection, the PCN- or CTX-untreated Δ*phpP* mutants and their wild-type strains were used as the controls.

### Detection of β-lactam antibiotic resistance of Δ*phpP* mutants

Susceptibility of the Δ*phpP*-6306, Δ*phpP*-SP5, Δ*phpP*-SP9 and Δ*phpP*-SP14 mutants to PCN or CTX was detected by E-test as described above. In the detection, the wild-type *S. pneumoniae* strains were used as the controls.

### Statistical analysis

Data from a minimum of three experiments were averaged and presented as mean ± standard deviation (SD). One-way analysis of variance (ANOVA) followed by Dunnett’s multiple comparisons test were used to determine significant differences. Statistical significance was defined as *p* <0.05.

## Results

### Increase of *stkP*- and *phpP*-mRNA levels induced by sublethal PCN and CTX

The *phpP*- and *stkP*-mRNA levels of each of the four tested *S. pneumoniae* strains were relatively lower. When the strains were treated with 1/4 MIC PCN or CTX, the *phpP*- and *stkP*-mRNA levels were rapidly elevated (Fig. 2A-2D). The data suggested that sublethal PCN and CTX can act as the stimulators to induce the expression of *S. pneumoniae phpP* and *stkP* genes.

**Fig. 2.**
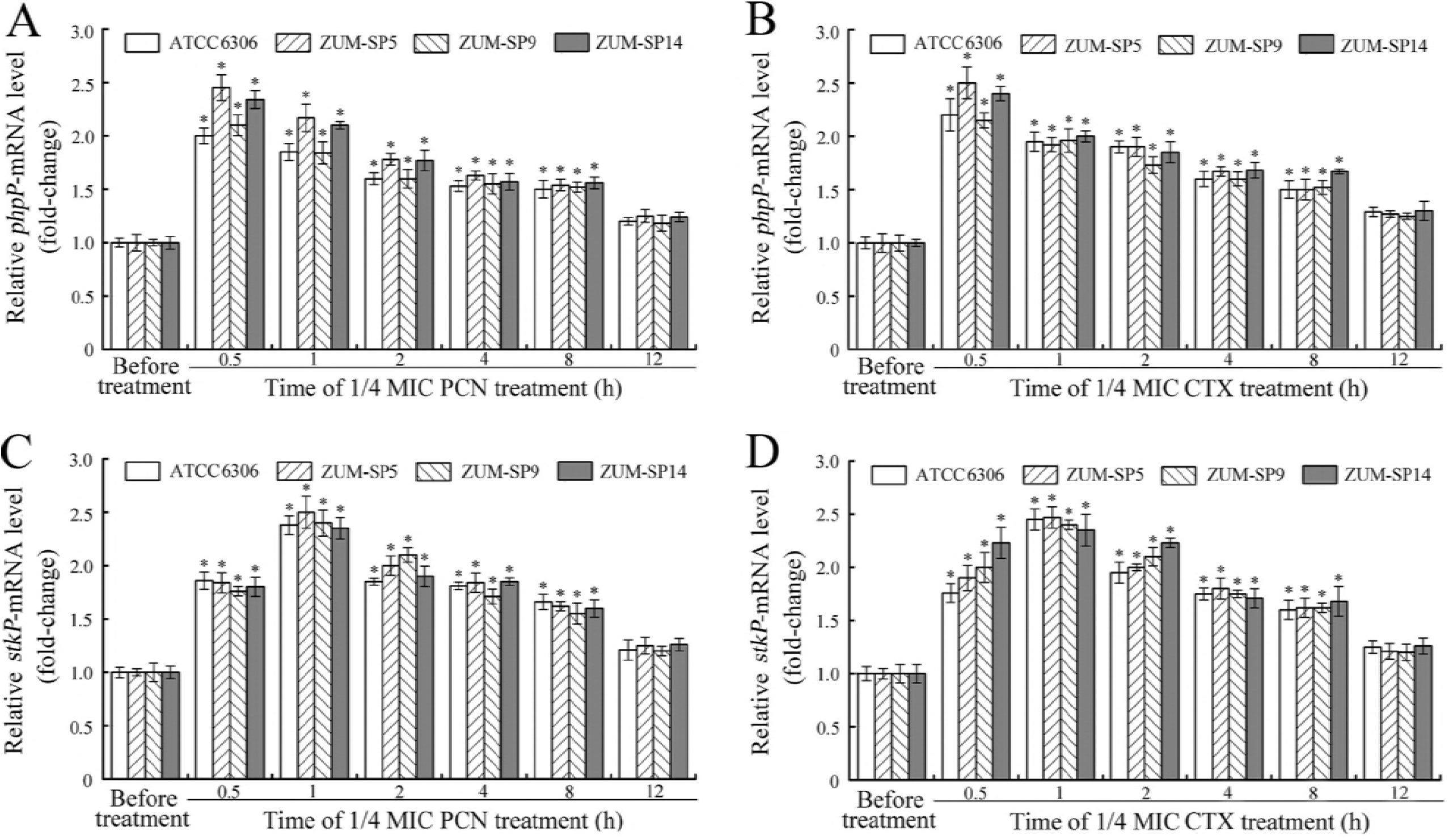
Increase of *phpP*- and *stkP*-mRNAs after treatment with sublethal PCN and CTX. (A). Increase of *phpP*-mRNA induced by 1/4 MIC PCN, detected by qRT-PCR. Bars show the mean ± SD of three independent experiments. The *phpP*-mRNA levels in the PCN-untreated *S. pneumoniae* strains (before treatment) were set as 1.0. *: *p*<0.05 *vs* the *phpP*-mRNA levels in the PCN-untreated strains. (B). Increase of *phpP*-mRNA induced by 1/4 MIC CTX, detected by qRT-PCR. The legend is the same as shown in A but for CTX-induced *phpP*-mRNA detection. (C). Increase of *stkP*-mRNA induced by 1/4 MIC PCN, detected by qRT-PCR. Bars show the mean ± SD of three independent experiments. The *stkP*-mRNA levels in the PCN-untreated *S. pneumoniae* strains (before treatment) were set as 1.0. *: *p*<0.05 *vs* the *stkP*-mRNA levels in the PCN-untreated strains. (B). Increase of *stkP*-mRNA induced by 1/4 MIC of CTX, detected by qRT-PCR. The legend is the same as shown in C but for CTX-induced *stkP*-mRNA detection.

### Characterization of *phpP* and *stkP* genes

The product of *S. pneumoniae phpP* or *stkP* gene was predicted as a secretary cytosolic or a transmembrane protein (Fig. 3A and 3B). The *phpP* gene contains a PP2Cc type protein phosphatase domain (6-238 aa) with five enzymatic active sites (Fig. 3C). The *stkP* gene contains a C type STK domain belonging to STKc_PknB superfamilay containing twelve ATP-binding sites, six dimer interface sites, two activation loops, eight polypeptide substrate-binding sites and sixteen enzymatic active sites as well as four penicillin-binding protein and STK-associated (PASTA) superfamily domains (Fig. 3D).

**Fig. 3.**
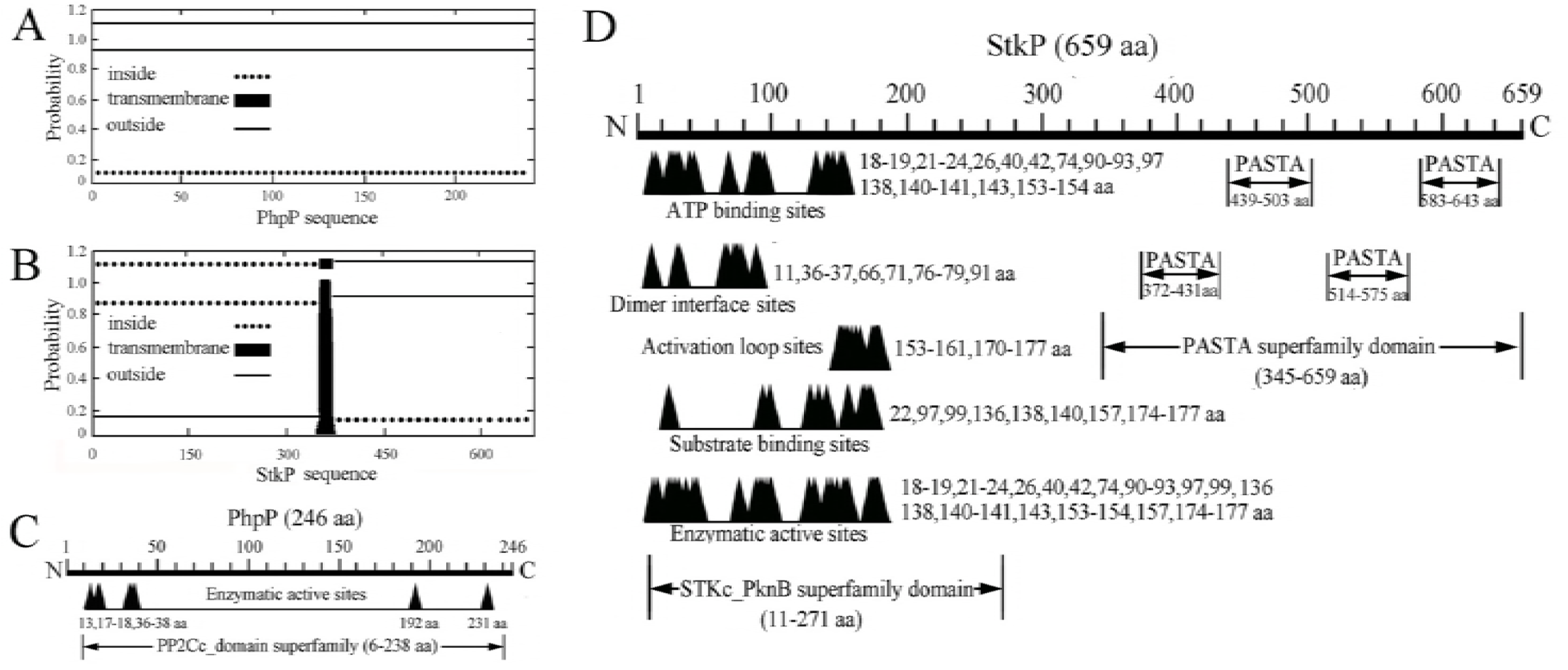
Predicted characteristics of *S. pneumoniae phpP* and *stkP* genes. (A). Structure and location of *phpP* gene product, predicted using TMHMM software. (B). Structure and location of *stkP* gene product, predicted using TMHMM software. (C). PP2C type phosphatase domain in *phpP* gene product (PhpP), predicted using NCBI Database CDD software. (D). STK and PASTA domains in *stkP* gene product (StkP), predicted using NCBI Database CDD software.

### Effects of expression and extraction of *stkP* and *phpP* genes

The nucleotide and amino acid sequence identities of *phpP* or *stkP* genes from the four *S. pneumoniae* strains were 99.3%-100% and 99.1%-100% or 99.2%-99.7% and 98.7%-99.9%, compared to the same two genes in GenBank (accession No.: NC003098) (data not shown). The two engineered strains expressed the target recombinant proteins rPhpP and rStkP, respectively, and the extracted rPhpP or rStkP was showed as a single band in gels (Fig. 4A-4D).

**Fig. 4.**
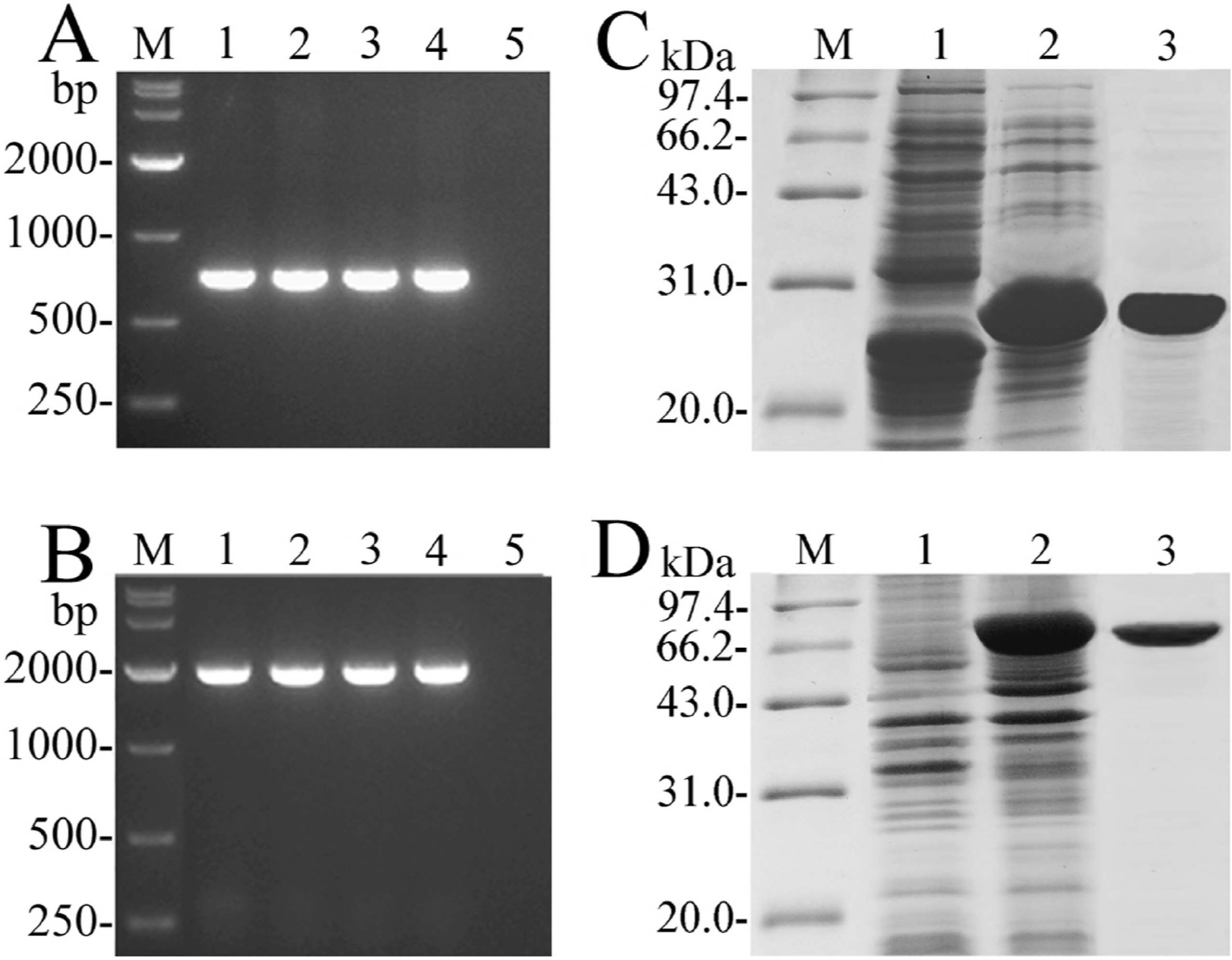
Amplification and expression of *S. pneumoniae phpP* and *stkP* genes. (A). Amplification fragments of *phpP* genes from *S. pneumoniae* strains, determined by PCR. Lane M: DNA marker. Lanes 1-4: amplicoms of *phpP* genes from *S. pneumoniae* strains ATCC6306, SP5, SP9 and SP14 (738 bp). Lane 5: blank control. (B). Amplification fragments of *stkP* genes from *S. pneumoniae* strains, determined by PCR. The legend is the same as shown in A but for *stkP* gene amplification (1977 bp). (C). Expression and extraction effects of rPhpP, detected by SDS-PAGE. Lane M: protein marker. Lane 1: wild-type *E. coli* BL21DE3. Lane 2: the expressed rPhpP (~28.3 kDa). Lane 3: the extracted rPhpP by Ni-NTA affinity chromatography. (D). Expression and extraction effects of rStkP, detected by SDS-PAGE. The legend is the same as shown in C but for rStkP expression and extraction (~75.8 kDa).

### Powerful Ser/Thr protein phosphatase activity of rPhpP

The rPhpP from *S. pneumoniae* ATCC6306 hydrolyzed p-NPP, an universal Ser/Thr phosphatase substrate, and RRA(pT)VA, a PP2C type Ser/Thr protein phosphatase-specific substrate, with concentration-dependent manners (Fig. 5A). The Km and Kcat values of rPhpP hydrolyzing RRA(pT)VA substrate were 277.35 μmol/L and 0.71 S^−1^, respectively (Fig. 5B). Moreover, the PP2C type Ser/Thr protein phosphatase inhibitor NaF, but not the PP1, PP2A or PP2B type Ser/Thr phosphatase inhibitor OA, inhibited the hydrolytic activity of rPhpP (Fig. 5C). The data suggested that the product of *S. pneumoniae phpP* gene is a PP2C type Ser/Thr protein phosphatase.

**Fig. 5.**
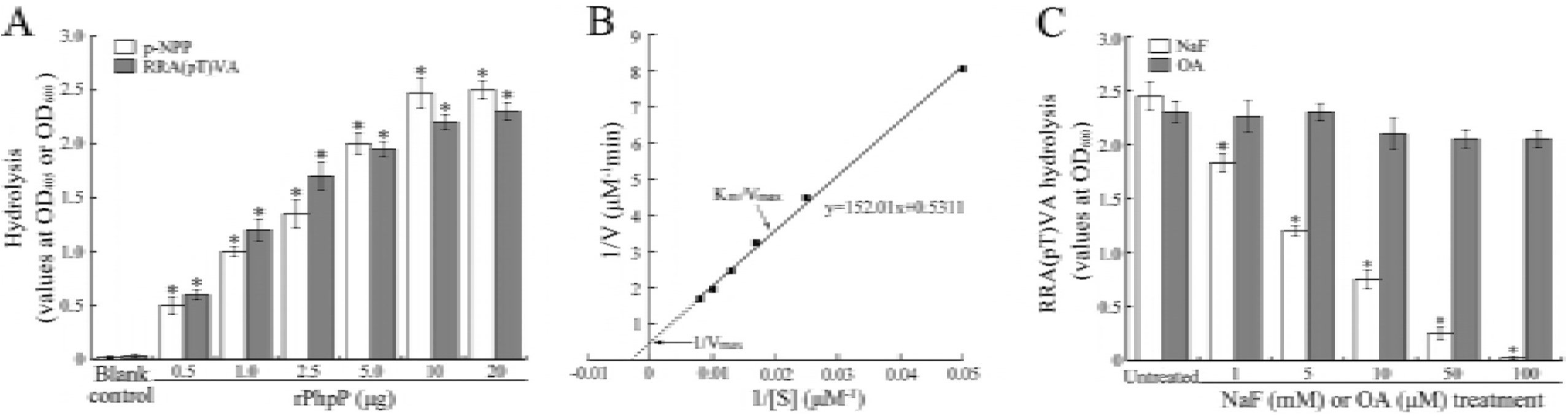
Powerful protein phosphatase activity of rPhpP from *S. pneumoniae*. (A). Ability of rPhpP hydrolyzing phosphatase substrates p-NPP and RRA(pT)VA, determined by spectrophotometry. Bars show the means ± SD of three independent experiments. p-NPP is a universal Ser/Thr phosphatase substrate while RRA(pT)VA is a PP2C type Ser/Thr protein phosphatase-specific substrate. (B). Km and Kcat values of rPhpP hydrolyzing RRA(pT)VA substrate, determined by spectrophotometry. 5 μg rPhpP and 50, 100, 150, 200 or 250 μM RRA(pT)VA phosphopeptide were used. (C). Enzymatic activity of rPhpP after treatment with phosphatase inhibitors, determined by spectrophotometry. NaF is a PP2C type Ser/Thr protein phosphatase inhibitor while OA is an inhibitor of PP1, PP2A or PP2B type Ser/Thr phosphatases.

### PCN/CTX-induced StkP phosphorylation and PhpP-caused StkP dephosphorylation

The Δ*phpP* mutants could grow persistently in the TH broth similarly to the wild-type strains (Fig. 6A). The PCR plus sequencing and Western Blot assay confirmed the *phpP* gene knockout and no PhpP expression in all the four Δ*phpP* mutants (Fig. 6B and 6C). 1/4 MIC PCN or CTX could cause the increase of StkP phosphorylation levels in the Δ*phpP* mutants and their wild-type strains (Fig. 6D and 6E). However, the Δ*phpP* mutants presented significantly higher PCN- or CTX-induced StkP phosphorylation levels than the wild-type strains (Fig. 6D and 6E). The data suggested that sublethal PCN and CTX can induce phosphorylation of StkP and PhpP can cause dephosphorylation of StkP *in vivo*.

**Fig. 6.**
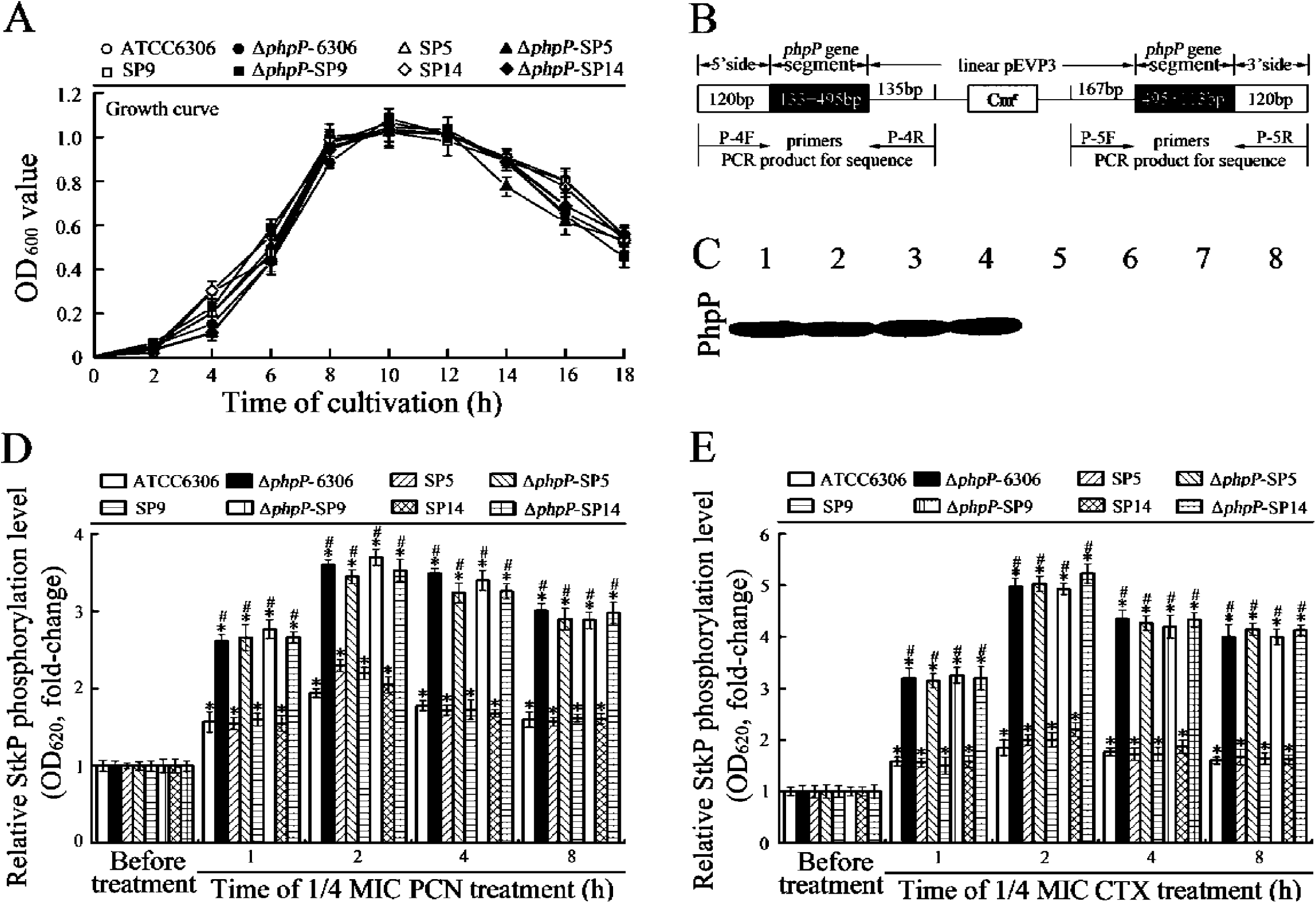
PCN/CTX-induced StkP phosphorylation and PhpP-caused StkP-dephosphorylation. (A). Growth curves of the Δ*phpP* mutants and wild-type strains in TH broth, determined by spectrophotometry. Bars show the means ± SD of three independent experiments. (B). Schematic diagram of PCR and sequencing results of the Δ*phpP* mutants. (C). PhpP absence in the Δ*phpP* mutants, determined by Western Blot assay. Lanes 1-4: immunoblotting bands of PhpP proteins from the wild-type *S. pneumoniae* strain ATCC 6306 and *S. pneumoniae* isolates SP1, SP5 and SP9. Lanes 5-8: no immunoblotting bands of PhpP proteins in the Δ*phpP* mutants. (D) Sublethal PCN-induced increase of StkP phosphorylation and decrease of StkP phosphorylation in the Δ*phpP* mutants, detected by spectrophotometry. Bars show the means ± SD of three independent experiments. *: *p*<0.05 *vs* the StkP phosphorylation levels of the PCN-untreated wild-type strains and Δ*phpP* mutants. ^#^: *p*<0.05 *vs* the StkP phosphorylation levels of the wild-type strains. (E) Sublethal CTX-induced increase of StkP phosphorylation and decrease of StkP phosphorylation in the Δ*phpP* mutants, detected by spectrophotometry. The legend is the same as shown in D but for detection of CTX-induced StkP phosphorylation.

### Increased PCN and CTX resistance of Δ*phpP* mutants

The MICs of PCN and CTX against wild-type ATCC6306, SP5, SP9 and SP14 strains of *S. pneumoniae* were 0.02-0.5 μg/ml, indicating all the strains were sensitive to both PCN and CTX. However, the MICs of PCN and CTX against the four Δ*phpP* mutants were significantly elevated as 4-16 µg/ml (Table 2). The data suggested that PhpP is involved in β-lactam antibiotic resistance of *S. pneumoniae*.

**Table 2.** MICs of PCN and CTX against *S. pneumoniae* strains.

| Strain | MIC (μg/ml) | |
| --- | --- | --- |
|  | PCN | CTX |
| Wild-type ATCC6306 | 0.05 | 0.12 |
| Δ*phpP*-6306 | 8 | 8 |
| Wild-type SP5 | 0.05 | 0.25 |
| Δ*phpP*-SP5 | 8 | 16 |
| Wild-type SP9 | 0.25 | 0.5 |
| Δ*phpP*-SP9 | 16 | 16 |
| Wild-type SP14 | 0.02 | 0.03 |
| Δ*phpP*-SP14 | 4 | 4 |

## Discussion

Kinases play key roles in signal transduction in both eukaryotes and prokaryotes and are activated by either phosphorylation or dephosphorylation [44]. Unlike eukaryotes, histidine kinases but not serine/threonine/tyrosine (Ser/Thr/Tyr) kinases act as the major signal sensors and transducers in prokaryotic bacteria [45]. Previous studies confirmed that StkP of *S. pneumoniae* is an eukaryotic-type Ser/Thr protein kinase containing PBP-like domains in its extracellular region and Ser/Thr phosphorylation sites in its intracellular region [46], and PhpP and StkP of *S. pneumoniae* compose a StkP-PhpP signaling couple [25]. Operon is a genic unit for transcription in prokaryotic genome that composed of promoter, operator and function-associated structural genes [47]. Our bioinformatic analysis showed that PhpP- and StkP-encoding genes of *S. pneumoniae* are located in the same one operon for co-expression, implying the close functional association between the two genes. As described in the introduction, StkP of *S. pneumoniae* is involved in PBP mutation-independent penicillin resistance [22]. Therefore, the product of *phpP* gene, PhpP, may act as a protein phosphotase to down-regulate StkP phosphorylation level by dephosphorylation to participate in StkP-involved β-lactam antibiotic resistance.

According to the differences of amino acid sequences and structures, Ser/Thr protein phosphatases are classified into MPP and PPP families as well as the former contains Mg^2+^ or Mn^2+^-dependent PP2C type phosphatases while the latter includes PP1, PP2A and PP2B type phosphatases [48]. PP2C type phosphatases has an extensive distribution in bacteria, yeasts, plants and mammalian cells to play various and complicated functions such as signaling transduction by dephosphorylation, cellular generation cycle regulation, monitoring DNA damage and RNA splicing [48-50]. Our bioinformatic prediction showed that *phpP* gene of *S. pneumoniae* contains a PP2C type protein phosphatase domain. The recombinant expression product of *phpP* gene (rPhpP) hydrolyzed both the general Ser/Thr phosphatase substrate p-NPP and PP2C type Ser/Thr protein phosphatase-specific substrate RRA(pT)VA in a concentration-dependent manner but its RRA(pT)VA-hydrolyzed ability was blocked by the PP2C type Ser/Thr protein phosphatase-specific inhibitor NaF. Previous studies confirmed that StkP and PhpP of *S. pneumoniae* compose a StkP/PhpP signaling couple in which StkP is activated by Ser/Thr residual phosphorylation and PhpP could hydrolyze phosphoryl groups in StkP *in vitro* [25,51]. However, we confirmed that the sublethal PCN and CTX induced the phosphorylation of StkP and the StkP phosphorylation levels in the Δ*phpP* mutants were significantly higher than that in the wild-type strains. All the data suggested that the product of *S. pneumoniae phpP* gene is a PP2C type Ser/Thr protein phosphatase to play a reverse regulation role in StkP/PhpP signaling couple by dephosphorylation of StkP.

Previous studies reported that penicillin-binding protein and STK-associated (PASTA) superfamily domains in some proteins from many Gram-positive bacteria have a potential ability to bind to β-lactam antibiotics [52,53], and our bioinformatic analysis also revealed that StkP of *S. pneumoniae* possesses four PASTA domains located its carboxyl terminal, implying that the StkP may served as the β-lactam antibiotic receptor and β-lactam antibiotics may cat as environmental activators of StkP-PhpP signaling couple. In the present study, the sublethal PCN or CTX (1/4 MIC) caused the significant increase of *stkP*- and *phpP*-mRNA levels and StkP phosphorylation. The data suggested that StkP-PhpP couple is a β-lactam antibiotic-associated signaling pathway of *S. pneumoniae*.

Bacterial PBP is a group of peptidoglycan biosynthesis-associated transpeptidases, carboxypeptidases and endopeptidases that located on surface of bacteria [54]. β-lactam antibiotics can bind to the peptidases to cause them inactivation by due to enzymatic molecular allosterism but PBP mutation can cause β-lactam antibiotic resistance due to the decrease of β-lactam antibiotic-binding ability of PBP [55,56]. However, some *S. pneumoniae* strains presented PBP mutation-independent β-lactam antibiotic resistance [20-22]. In the present study, all the tested *S. pneumoniae* strains were sensitive to both PCN and CTX (MICs: 0.02-0.5 μg/ml). When the *phpP* genes were knockout, the Δ*phpP* mutants became resistant to the two antibiotics (MICs: 4-16 μg/ml). The data imply that StkP phosphorylation promotes but PhpP-based StkP dephosphorylation inhibits the generation of PBP mutation-independent β-lactam antibiotic resistance.

## Conclusions

Sublethal PCN and CTX can act as environmental inducers to cause the increase of *stkP* and *phpP* gene expression and StkP phosphorylation of *S. pneumoniae*. The product of *phpP* gene is a PP2C type Ser/Thr protein phosphatase to cause dephosphorylation of StkP that plays a reverse regulating role in StkP/PhpP signaling couple. Sublethal β-lactam antibiotics can act as environmental inducers of *phpP* and *stkP* gene expression and knockout of *phpP* gene caused the significant increase of β-lactam antibiotic resistance of *S. pneumoniae*.

## Additional file

This manuscript has no additional files.

## Abbreviations

*S. pneumoniae*: *Streptococcus pneumoniae*
PNC: penicillin
CTX: cefotaxime
qRT-PCR: quantitative reverse transcription polymerase chain reaction
Δ*phpP*: *phpP* gene-knockout mutant
MIC: minimal inhibitory concentration
mRNA: messenger ribonucleic acid
PBP: penicillin-binding proteins
STK: serine/threonine kinase
DNA: deoxyribonucleic acid
TH: Todd-Hewitt
LB: Luria-Bertani
OD: optical density
PBS: phosphate buffered saline
RNA: ribonucleic acid
SDS-PAGE: sodium dodecyl sulfate polyacrylamide gel electropheresis
CDD: conserved domain database
IPTG: isopropy-β-D-thiogalactoside
OA: Okadaic acid
NaF: sodium fluoride
SD: standard deviation
PASTA: penicillin-binding protein and STK-associated
TB: Tris-HCl buffer

## Acknowledgements

We are grateful to Dr. Morrisson (University of Illinois, Chicago, USA) for kindly providing the pEVP3 plasmid used in this study.

## Availability of data and materials

The datasets analyzed during the current study are available from the corresponding author on reasonable request.

## Author’s contributions

A.H.S. and J.Y. conceived and designed the experiments. Y.Y.H., Y.H.S. and P.D. performed the experiments. Y.Y.H., Y.H.S. and X.X.L. analyzed the data. Y.Y.H., A.H.S. and J.Y. wrote the manuscript. A.H.S. obtained the funding. All authors have read and approved the manuscript.

## Ethics approval and consent to participate

This study has no medical ethic problems. All the animal experimental protocols were approved by the Ethics Committee for Animal Experiment of Hangzhou Medical College.

## Consent for publication

Not applicable.

## References

1. O’Brien KL, Wolfson LJ, Watt JP, Henkle E, Deloria-Knoll M, McCall N, Lee E, Mulholland K, Levine OS, Cherian T; Hib and Pneumococcal Global Burden of Disease Study Team. Burden of disease caused by *Streptococcus pneumoniae* in children younger than 5 years: global estimates. Lancet, 2009;374:893–902.

2. Yao KH, Yang YH. *Streptococcus pneumoniae* diseases in Chinese children: past, present and future. Vaccine, 2008;26(35):4425–4433.

3. Henriques-Normark B, Tuomanen EI. The pneumococcus: Epidemiology, microbiology, and pathogenesis. Cold Spring Harb Perspect Med, 2013;3:a010215–010229.

4. Leimkugel J, Forgor AA, Gagneux S, Pflüger V, Flierl C, Awine E, Naegeli Mn, Dangy JP, Smith T, Hodgson A, Pluschke G. An outbreak of serotype 1 *Streptococcus pneumoniae* meningitis in Northern Ghana with features that are characteristic of *Neisseria meningitidis* meningitis epidemics. J Infect Dis, 2005;15:192–199.

5. Chen Y, Deng W, Wang SM, Mo QM, Jia H, Wang Q, Li SG, Li X, Yao BD, Liu CJ, Zhan YQ, Ji C, Lopez AL, Wang XY. Burden of pneumonia and meningitis caused by *Streptococcus pneumoniae* in China among children under 5 years of age: A systematic literature review. PLoS One, 2011;6(11):e27333–27341.

6. Iovino F, Orihuela CJ, Moorlag HE, Molema G, Bijlsma JJE. Interactions between blood-borne *Streptococcus pneumoniae* and the blood-brain barrier preceding meningitis. PLoS One, 2013;8(7):e68408–68420.

7. Sfaihi L, Kamoun F, Kamoun T, Aloulou H, Mezghani S, Hammemi A, Hachicha M. Bacterial meningitis in children: epidemiological data and outcome. Tunis Med, 2014;92(2):141–146.

8. Brink M, Welinder-Olsson C, Hagberg L. Time window for positive cerebrospinal fluid broad-range bacterial PCR and *Streptococcus pneumoniae* immunochromatographic test in acute bacterial meningitis. Infect Dis, 2015;47:869–877.

9. Felmingham D, Cantón R, Jenkins SG. Regional trends in beta-lactam, macrolide, fluoroquinolone and telithromycin resistance among *S. pneumoniae* isolates 2001-2004. J Infect, 2007;55(2):111–118.

10. Song JH, Jung SI, Ko KS. High prevalence of antimicrobial resistance among clinical *Streptococcus pneumoniae* isolates in Asia. Antimicrob Agents Chemother, 2004; 48:2101–2107.

11. Yang F, Xu XG, Yang MJ, Zhang YY, Klugman KP, McGee L. Antimicrobial susceptibility and molecular epidemiology of *Streptococcus pneumoniae* isolated from Shanghai, China. Int J Antimicrob Agents, 2008;32(5):386–391.

12. Kumarasamy KK, Toleman MA, Walsh TR, et al. Emergence of a new antibiotic resistance mechanism in India, Pakistan, and the UK: a molecular, biological, and epidemiological study. Lancet Infect Dis, 2010;10:597–602.

13. Hackel M, Lascols C, Bouchillon S, Hilton B, Morgenstern D, Purdy J. Serotype prevalence and antibiotic resistance in *Streptococcus pneumoniae* clinical isolates among global populations. Vaccine, 2013;31(42):4881–4887.

14. Moujaber GE, Osman M, Rafei R, Dabboussi F, Hamze M. Molecular mechanisms and epidemiology of resistance in *Streptococcus pneumoniae* in the Middle East region. J Med Microbiol, 2017;66:847–858.

15. Jacobs MR. Antimicrobial-resistant *Streptococcus pneumoniae*: trends and management. Expert Rev Anti Infect Ther, 2008;6(5):619–635.

16. Huang S, Liu X, Lao W, Zeng S, Liang H, Zhong R, Dai X, Wu X, Li H, Yao Y. Serotype distribution and antibiotic resistance of *Streptococcus pneumoniae* isolates collected at a Chinese hospital from 2011 to 2013. BMC Infect Dis, 2015;15(1):312–322.

17. Amin AN, Cerceo EA, Deitelzweig SB, Pile JC, Rosenberg DJ, Sherman BM. The hospitalist perspective on treatment of community-acquired bacterial pneumonia. Postgrad Med, 2014;126(2)18–29.

18. Contreras-Martel C, Dahout-Gonzalez C, Martins ADS, Ados S, Kotnik M, Dessen A. PBP active site flexibility as the key mechanism for β-lactam resistance in pneumococci. J Mol Biol, 2009;387(4):899–909.

19. Albarracin-Orio AG, Pinas GE, Cortes PR, et al. Compensatory evolution of pbp mutations restores the fitness cost imposed by beta-lactam resistance in *Streptococcus pneumoniae*. PLoS Pathog, 2011;7(2):e1002000–1002033.

20. Chesnel L, Carapito R, Croizé J, Dideberg O, Vernet T, Zapun A. Identical penicillin-binding domains in penicillin-binding proteins of *Streptococcus pneumoniae* clinical isolates with different levels of beta-lactam resistance. Antimicrob Agents Chemother. 2005;49(7):2895–2902.

21. Hakenbeck R, Brückner R, Denapaite D, Maurer P. Molecular mechanisms of β-lactam resistance in *Streptococcus pneumonia*. Future Microbiol, 2012;7(3):395–410.

22. Dias R, Félix D, Canica M, Trombe MC. The highly conserved serine threonine kinase StkP of *Streptococcus pneumoniae* contributes to penicillin susceptibility independently from genes encoding penicillin-binding proteins. BMC Microbiol, 2009;9(5):121–129.

23. Tettelin H, Nelson KE, Paulsen IT, Eisen JA, Read TD, Peterson S, Heidelberg J, DeBoy RT, Haft DH, Dodson RJ, Durkin AS, Gwinn M, Kolonay JF, Nelson WC, Peterson JD, Umayam LA, White O, Salzberg SL, Lewis MR, Radune D, Holtzapple E, Khouri H, Wolf AM, Utterback TR, Hansen CL, McDonald LA, Feldblyum TV, Angiuoli S, Dickinson T, Hickey EK, Holt IE, Loftus BJ, Yang F, Smith HO, Venter JC, Dougherty BA, Morrison DA, Hollingshead SK, Fraser CM. Complete genome sequence of a virulent isolate of *Streptococcus pneumoniae*. Science, 2001;293(5529):498–506.

24. Hoskins JA, Alborn W Jr., Arnold J, Blaszczak L, Burgett S, DeHoff BS, Estrem S, Fritz L, Fu DJ, Fuller W, Geringer C, Gilmour R, Glass JS, Khoja H, Kraft A, LaGace R, LeBlanc DJ, Lee LN, Lefkowitz EJ, Lu J, Matsushima P, McAhren S, McHenney M, McLeaster K, Mundy C, Nicas TI, Norris FH, O’Gara M, Peery R, Robertson GT, Rockey P, Sun PM, Winkler ME, Yang Y, Young-Bellido M, Zhao G, Zook C, Baltz RH, Jaskunas S. Richard, Rosteck PR Jr, Skatrud PL, Glass JI. Genome of the bacterium *Streptococcus pneumoniae* strain R6. J Bacteriol, 2001;183 (19):5709–5717.

25. Osaki M, Arcondéguy T, Bastide A, Touriol C, Prats H, Trombe MC. The StkP/PhpP signaling couple in *Streptococcus pneumoniae*: cellular organization and physiological characterization. J Bacteriol, 2009;191(15):4943–4950.

26. Beier D, Gross R. Regulation of bacterial virulence by two-component systems. Curr Opin Microbiol, 2006, 9(2):143–152.

27. Rogers PD, Liu TT, Barker KS, Hilliard GM, English BK, Thornton J, Swiatlo E, McDaniel LS. Gene expression profiling of the response of *Streptococcus pneumoniae* to penicillin. J Antimicrob Chemother, 2007;59(4):616–626.

28. Hesje CK, Drlica K, Blondeau JM. Mutant prevention concentration of tigecycline for clinical isolates of Streptococcus pneumoniae and Staphylococcus aureus. J Antimicrob Chemother, 2015;70:494–497.

29. National Committee for Clinical Laboratory Standards (NCCLS). Performance standards for antimicrobial susceptibility testing. 25th informational supplement (M100-S25), Wayne: NCCLS, PA, USA, 2015.

30. Zhang L, Zhang CL, Ojcius DM, Sun D, Zhao JF, Lin XA, Li LW, Li LJ, Yan J. The mammalian cell entry (Mce) protein of pathogenic *Leptospira* species is responsible for RGD motif-dependent infection of cells and animals. Mol Microbiol, 2012;83(5):1006–1023.

31. El Aila NA, Emler S, Kaijalainen T, De Baere T, Saerens B, Alkan E, Deschaght P, Verhelst R, Vaneechoutte M. The development of a 16S rRNA gene based PCR for the identification of *Streptococcus pneumoniae* and comparison with four other species specific PCR assays. BMC Infect Dis, 2010;10:104–112.

32. Pfaffl MW, Horgan GW, Dempfle L. Relative expression software tool (REST) for group-wise comparison and statistical analysis of relative expression results in real-time PCR. Nucleic Acids Res, 2002;30(9):e36–46.

33. Marchler-Bauer A, Lu S, Anderson JB, Chitsaz F, Derbyshire MK, DeWeese-Scott C, Fong JH, Geer LY, Geer RC, Gonzales NR, Gwadz M, Hurwitz DI, Jackson JD, Ke Z, Lanczycki CJ, Lu F, Marchler GH, Mullokandov M, Omelchenko MV, Robertson CL, Song JS, Thanki N, Yamashita RA, Zhang D, Zhang N, Zheng C, Bryant SH. CDD: a Conserved Domain Database for the functional annotation of proteins. Nucleic Acids Res, 2011;39:D225–D229.

34. Wang H, Wu YF, Ojcius DM, Yang XF, Zhang CL, Ding SB, Lin XA, Yan J. Leptospiral hemolysins induce proinflammatory cytokines through Toll-like receptor 2- and 4-mediated JNK and NF-κB signaling pathways. PLoS One, 2012;7(8):e42266–42280.

35. Lee, SW, Jeong KS, Han SW, Lee SE, Phee BK, Hahn TR, Ronald P. The *Xanthomonas oryzae* pv. oryzae PhoPQ two-component system is required for AvrXA21 activity, *hrpG* expression, and virulence. J Bacteriol, 2008;190(6):2183–2197.

36. Treuner-Lange A, Ward MJ, Zusman DR. Pph1 from *Myxococcus xanthus* is a protein phosphatase involved in vegetative growth and development. Mol Microbiol, 2001; 40(1):126–140.

37. Garcia L, Garcia F, Llorens F, Unzeta M, Itarte E, Gómez N. PP1/PP2A phosphatases inhibitors okadaic acid and calyculin A block ERK5 activation by growth factors and oxidative stress. FEBS Lett, 2002;523(1-3):90–94.

38. Marley AE, Sullivan JE, Carling D, Abbott WM, Smith GJ, Taylor IW, Carey F, Beri RK. Biochemical characterization and deletion analysis of recombinant human protein phosphatase 2C. Biochem J, 1996;320(Pt3):801–806.

39. Kassegne K, Hu WL, Ojcius DM, Sun D, Ge YM, Zhao JF, Yang XF, Li LJ, Yan J. Identification of collagenase as a critical virulence factor for invasiveness and transmission of pathogenic *Leptospira species*. J Infect Dis, 2014;209(7):1105–1115.

40. Pestova EV, Morrison DA. Isolation and characterization of three *Streptococcus pneumoniae* transformation-specific loci by use of a *lacZ* reporter insertion vector. J Bacteriol, 1998;180(10):2701–2710.

41. Thanassi JA, Hartman-Neumann SL, Dougherty TJ, Dougherty BA, Pucci MJ. Identification of 113 conserved essential genes using a high-throughput gene disruption system in *Streptococcus pneumoniae*. Nucleic Acids Res, 2002;30(14):3152–3162.

42. Contreras-Martel C, Dahout-Gonzalez C, Martins-Ados S, Kotnik M, Dessen A. PBP active site flexibility as the key mechanism for beta-lactam resistance in pneumococci. J Mol Biol, 2009;387(4):899–909.

43. Fruehling S, Longnecker R. In *vitro* assays for the detection of protein tyrosine phosphorylation and protein tyrosine kinase activities. Methods Mol Biol, 2001; 174:337–343.

44. Kumar R, Deivendran S, Santhoshkumar TR, Pillai MR. Signaling coupled epigenomic regulation of gene expression. Oncogene, 2017;36(43):5917–5926.

45. Khorchid A, Mkura M. Bacterial histidine kinase as signal sensor and transducer. Int J Biochem Cell Biol, 2006;38(3):307–312.

46. Giefing C, Jelencsics KE, Gelbmann D, Senn BM, Nagy E. The pneumococcal eukaryotic-type serine/threonine protein kinase StkP co-localizes with the cell division apparatus and interacts with FtsZ *in vitro*. Microbiology, 2010;156(6):1697–1707.

47. Cherry L. Genome size and operon content. J Theror Biol, 2003;221(3):401–410.

48. Das AK, Helps NR, Cohen PT, Barford D. Crystal structure of the protein serine/threonine phosphatase 2C at 2.0 A resolution. EMBO J, 1996;15 (24):6798–6809.

49. Schweighofer A, Hirt H, Meskiene I. Plant PP2C phosphatases: emerging functions in stress signaling. Trends Plant Sci, 2004;9(5):236–243.

50. Rajagopal L, Clancy A, Rubens CE. A eukaryotic type serine/threonine kinase and phosphatase in *Streptococcus agalactiae* reversibly phosphorylate an inorganic pyrophosphatase and affect growth, cell segregation, and virulence. Biol Chem J, 2003; 278(16):14429–14441.

51. Nováková L, Sasková L, Pallová P, Janeček J, Novotná J, Ulrych A, Echenique J, Trombe MC, Branny P. Characterization of a eukaryotic type serine/threonine protein kinase and protein phosphatase of *Streptococcus pneumoniae* and identification of kinase substrates. FEBS J, 2005;272(5):1243–1254.

52. Yeats C, Finn RD, Bateman A. The PASTA domain: a β-lactam-binding domain. Trends Biochem Sci, 2002;27(9):438–440.

53. Maestro B, Nováková L, Hesek D, Lee M, Leyva E, Mobashery S, Sanz JM, Branny P. Recognition of peptidoglycan and β-lactam antibiotics by extracellular domain of the Ser/Thr protein kinase StkP from *Streptococcus pneumonia*. FEBS Lett, 2011; 585(2):357–363.

54. Sauvage E, Kerff F, Terrak M, Ayala JA, Charlier P. The penicillin-binding proteins: Structure and role in peptidoglycan biosynthesis. FEMS Microbiol Rev, 2008;32:234–258.

55. Macheboeuf P, Contreras-Martel C, Job V, Dideberg O, Dessen A. Penicillin binding proteins: Key players in bacterial cell cycle and drug resistance processes. FEMS Microbiol Rev, 2006;30:673–691.

56. Zapun A, Contreras-Martel C, Vernet T. Penicillin-binding proteins and β-lactam resistance. FEMS Microbiol Rev, 2008;32:361–385.

